# NanoPAMDA enables rapid PAM characterization of metagenome-derived Cas9 effectors

**DOI:** 10.64898/2026.11.04.764188

**Authors:** Ibrahim Genel, Ceren Suren, Ibrahim C. Kurt

## Abstract

CRISPR-Cas9 orthologs remain functionally underexplored despite the rapidly growing scale of publicly available metagenomic sequencing data. Here, we applied a multi-stage computational discovery pipeline to two independent sources of metagenomic sequence data, a multi-biome collection of 620,185 metagenome-assembled genomes spanning 17 public catalogs, and raw sequencing reads from human microbiome studies assembled de novo, to identify novel Cas9 orthologs. Candidates were filtered for guide RNA architectural plausibility, sequence novelty, conservation of catalytic residues, and structural integrity, yielding 14 candidates spanning both data sources for experimental characterization. To determine PAM preference at scale, we developed a cost-effective Nanopore long-read-based adaptation of the high-throughput PAM determination assay (NanoPAMDA), enabling rapid, time-resolved, kinetic PAM profiling across many candidates in parallel. One Cas9 effector, E10, showed PAM-specific activity in vitro and was catalytically active in *E. coli*. Together, these findings demonstrate a scalable, long-read sequencing-based framework for discovering and biochemically characterizing novel Cas9 orthologs from metagenomic data.

## INTRODUCTION

The discovery that the CRISPR-Cas9 system could be repurposed as a programmable, RNA-guided DNA endonuclease transformed genome editing across biology and medicine. Following the initial biochemical characterization of the *Streptococcus pyogenes* Cas9 (SpCas9) dual-RNA (crRNA:tracrRNA) mechanism **(Jinek et al., 2012)**, rapid demonstration of its function in mammalian and other eukaryotic cells **(Cong et al., 2013; Mali et al., 2013)** established CRISPR-Cas9 as the dominant platform for targeted genome engineering (Doudna & Charpentier, 2014).

Despite this success, SpCas9 has intrinsic constraints that limit its universal applicability. Target site selection is restricted to sequences immediately adjacent to an 5’-NGG-3’ protospacer-adjacent motif (PAM) **(Nishimasu et al., 2014)**, and the large size of the SpCas9 complicates packaging into size-limited delivery vehicles such as adeno-associated virus (AAV) **(Ran et al., 2015)**. These limitations motivated efforts to identify alternative, naturally occurring Cas9 orthologs with a more compact size and distinct PAM specificities. Characterization of Cas9 from *Staphylococcus aureus* demonstrated that a smaller ortholog could support efficient genome editing and single-AAV-vector delivery **(Ran et al., 2015)**. Subsequent efforts identified additional compact orthologs, including Cas9 from *Campylobacter jejuni* **(Kim et al., 2017)**, and Cas9 from *Neisseria meningitidis* **(Edraki et al., 2019)**. Each of these orthologs recognizes its own distinct PAM **(Ran et al., 2015; Kim et al., 2017; Edraki et al., 2019)**. Therefore, naturally occurring Cas9 diversity is useful for expanding the accessible target space and delivery options. However, each newly characterized ortholog must be individually, and often laboriously, validated for PAM preference and editing activity. Reconstruction of the evolutionary pathway connecting IscB and TnpB nucleases, ancestors of Cas9 and Cas12, respectively, to modern CRISPR-Cas effectors demonstrated that RNA-guided DNA cleavage activity is a broadly distributed property across a continuum of related nuclease families found throughout bacterial and archaeal genomes **(Altae-Tran et al., 2021)**. This evolutionary continuity suggests that substantial functional diversity in Cas9 likely remains uncharacterized in existing and rapidly growing genomic databases.

The scale of available microbial genomic data has grown exponentially in the past decade, driven largely by metagenome-assembled genomes (MAGs). Large-scale surveys such as the Genomes from Earth’s Microbiomes (GEM) catalog (**Nayfach et al., 2021)** and expansive human microbiome sequencing efforts **(Pasolli et al., 2019)** have contributed MAGs spanning uncultivated microbial lineages. This resulted in systematic computational efforts to mine Cas9 diversity directly from large metagenomic collections. Such efforts have ranged from biochemical characterization of a curated panel of Cas9 orthologs **(Gasiunas et al., 2020)**, to large-scale mining and PAM prediction across thousands of candidate Cas9 sequences **(Fang et al., 2026)**.

Fully leveraging this scale of genomic diversity for functional discovery requires computational methods that filter large candidate pools into a tractable subset for experimental characterization, since exhaustive biochemical validation of every candidate remains infeasible. Recent advances in protein structure prediction and comparison tools, including ESMFold **(Lin et al., 2023)**, Foldseek **(van Kempen et al., 2024)**, and protein-language-model-based PAM prediction **(Nayfach et al., 2026)**, now make it possible to computationally assess candidate novelty, structural integrity, and likely PAM preference prior to committing to labor-intensive experimental validation.

Here, we applied a computational discovery pipeline combined with direct biochemical characterization. Candidates were mined from a multi-biome collection of metagenome-assembled genomes and from raw sequencing reads of controlled-access human microbiome studies, filtered through successive criteria for guide RNA architectural plausibility, sequence novelty, catalytic residue conservation, and structural integrity, and characterized experimentally using a Nanopore-adapted implementation of the high-throughput PAM determination assay (Walton et al., 2020; Walton et al., 2021). Among the 14 candidates, E10, a Cas9 from a mouse gut metagenome, displayed an NGG-like PAM preference in vitro and was catalytically active in E. coli.

## RESULTS

### Discovery of novel Cas9 orthologs across two distinct metagenomic data sources

Our discovery pipeline **(Fig. 1A)** was applied in parallel to two distinct data sources: an environmental microbiome collection of 620,185 metagenome-assembled genomes (MAG) spanning 17 public catalogs **(Supplementary Table 1)**, and raw shotgun sequencing reads from two controlled-access human microbiome studies (HMP-A and TOPDECC) **(Supplementary Table 2)** assembled de novo with MetaHipMer2. This dual-source design captures both the breadth of publicly curated genomic diversity and orthologs that may be underrepresented in existing catalogs due to assembly or binning limitations.

**Figure 1.**
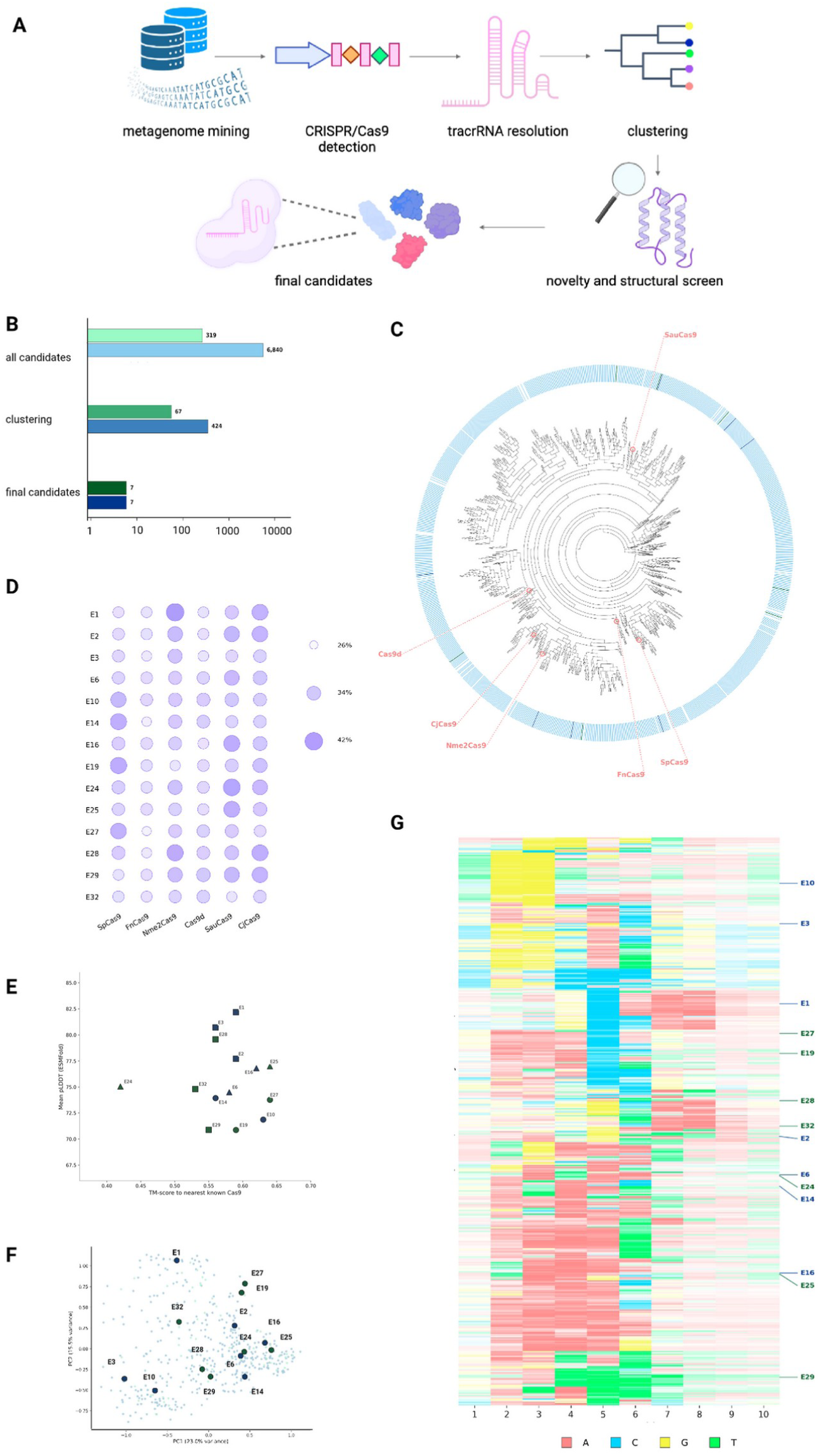
Metagenomic discovery pipeline and sequence diversity of mined Cas9 candidates. **(A)** Schematic of the computational discovery pipeline. Candidate genomic sequences were mined from metagenome-assembled genome catalogs and raw sequencing datasets (metagenome mining), screened for CRISPR arrays and adjacent Cas9-family genes (CRISPR/Cas9 detection), resolved for tracrRNA architecture (tracrRNA resolution), clustered to remove redundancy (clustering), screened for sequence novelty and structural integrity (novelty and structural screen), and narrowed to a final selected candidate set (final candidates). **(B)** Number of candidates retained at each stage of the discovery pipeline (log scale), shown separately for the environmental (clustered at 50% identity) and human microbiome (clustered at 90% identity) candidate pools, each ending in 7 candidates selected for experimental characterization. **(C)** Cladogram of sequence diversity across mined Cas9 candidates, constructed from cluster representatives spanning the combined candidate pool. The outer ring indicates the candidate source (environmental, light blue).Known Cas9 orthologs (SpCas9, FnCas9, Nme2Cas9, Cas9d, SauCas9, CjCas9) are indicated with callouts. **(D)** Pairwise amino acid sequence identity between each of the 14 selected candidates and six known Cas9 orthologs. **(E)** Mean per-residue confidence plotted against TM-score to the nearest known Cas9 ortholog, for all 14 selected candidates. Marker shape indicates each candidate’s closest structural match among the six known Cas9 orthologs used as comparators: circle, SpCas9; square, Nme2Cas9; triangle, SauCas9. Marker color indicates candidate source (navy, environmental; dark green, human microbiome). **(F)** Principal component projection of Protein2PAM-predicted PAM probability matrices across the discovered candidate pool (pale points, background) and the 14 selected candidates (labeled points). **(G)** Heatmap of predicted PAM identity across all discovered candidate representatives, hierarchically clustered by predicted PAM probability similarity. Each row is one candidate; columns correspond to the 10 predicted PAM positions. Cell color indicates the predicted nucleotide (A, red; C, blue; G, yellow; T, green), with color intensity scaled to prediction confidence. The 14 selected candidates are labeled on the right.

Screening the environmental metagenomics dataset for CRISPR arrays and adjacent Cas9-family proteins, resolving dual-guide RNA architecture, and filtering for tracrRNAs of 70–150 nucleotides identified 6,840 CRISPR systems **(Supplementary Table 3)**. This length window excludes truncated fragments and spuriously extended predictions arising from imperfect terminator detection. Clustering at 50% amino acid identity collapsed this set to 424 representatives **(Fig. 1B) (Supplementary Table 4)**. BLASTp screening against the NCBI non-redundant database (retaining only candidates with <90% identity to any characterized protein), combined with NUPACK-verified hairpin, nexus, and nexus-junction architectures, narrowed the pool to 51 sequences. A final round of catalytic and structural validation, assessing conservation across a six-reference Cas9 panel, domain architecture integrity, and ESMFold/Foldseek structural similarity, excluded two candidates for missing catalytic domains and four for substitutions at conserved catalytic residues, leaving 45. Seven of these were prioritized for experimental validation and downstream characterization by NanoPAMDA **(Supplementary Table 7)**.

A parallel pipeline was applied to the human microbiome metagenomic dataset. Of the 865 initial CRISPR systems, 319 passed the same-length filter **(Supplementary Table 5**). Given the smaller starting pool, clustering used a stricter 90% identity threshold, yielding 67 representatives **(Fig. 1B) (Supplementary Table 6)**. This arm was not subjected to BLASTp screening, gRNA structural filtering, or structural similarity assessment; only catalytic residue conservation was evaluated, and candidates were screened sequentially until seven satisfied the filter. In total, 14 candidates, seven from each source, were carried forward for cloning and NanoPAMDA-based PAM characterization.

### Sequence diversity and candidate distribution across the discovery pipeline

To situate the selected candidates within the diversity captured by the pipeline, we built a phylogenetic tree from 1,125 sequences: 1,105 non-redundant cluster representatives from re-clustering the combined pools at a looser 90% identity threshold, the 14 selected candidates, and six known Cas9 orthologs **(Fig. 1C)**. Pairwise amino acid identity between each candidate and the six known orthologs ranged from 26% to 42% **(Fig. 1D)**. The 14 candidates were distributed broadly across the tree rather than confined to a single clade, indicating that our screening criteria selected across the pipeline’s discovered diversity rather than from one phylogenetic corner of it. We next compared the predicted structure of each of the 14 candidates against the six known Cas9 orthologs, plotting structural confidence against similarity to the nearest match **(Fig. 1E)**. Six candidates (43%) were structurally closest to Nme2Cas9, four (29%) to SpCas9, and four (29%) to SauCas9, spanning both environmental and human microbiome candidates. TM-scores ranged from 0.42 (E24) to 0.64 (E25 and E27) **(Supplementary Table 9)**, indicating that while most candidates showed moderate structural similarity to a known ortholog, this similarity was not uniform across the panel.

### Predicted PAM diversity resolves into distinct sequence families

To assess PAM diversity across the 14-candidate Cas9 effector pool, we predicted in silico PAM preferences for all 491 candidate representatives using Protein2PAM. A principal component projection of the resulting predicted PAM probability profiles showed the 14 candidates distributed broadly across the landscape of discovered diversity rather than confined to one region **(Fig. 1F)**. Hierarchical clustering of the same per-position nucleotide calls resolved several visually distinct sequence families rather than a continuum **(Fig. 1G)**, indicating that predicted PAM diversity reflects real underlying sequence structure rather than prediction noise. A guanine-rich family at positions 2 to 3 included E10. A cytosine-dominant family centered on position 5 included E1, a broad adenine-rich family spanned much of the pool, and a thymine-rich family at positions 4 to 6 included E29. The 14 candidates carried forward for NanoPAMDA characterization spanned at least four of these families, consistent with the broader spread already observed by principal component analysis.

### NanoPAMDA enables rapid quantitative PAM profiling *in vitro* by Nanopore sequencing

To enable rapid experimental PAM profiling across the candidate panel, we adapted the high-throughput PAM determination assay (HT-PAMDA) to Oxford Nanopore sequencing and termed the resulting workflow NanoPAMDA **(Fig. 2A)**. NanoPAMDA preserves the time-resolved library-depletion principle of HT-PAMDA while coupling a single-step barcoded amplification strategy with Nanopore sequencing and custom computational demultiplexing to recover PAM-specific cleavage kinetics. We benchmarked the workflow using SpCas9, for which NanoPAMDA reproducibly recovered the canonical 5’-NGG-3’ PAM **(Fig. 2B)**. Across the three independent sequencing runs, 63.4–72.7% of reads were successfully demultiplexed per run **(Fig. 2C)**, and sequencing from library loading to basecalled output was completed within 3–4 h. Read depth across the 14 metagenome-derived Cas9 candidates was similarly consistent across timepoints for most of the panel **(Fig. 2C)**, with one exception: E1 was recovered at markedly lower and more slowly declining depth than the other 13 candidates. Having established that NanoPAMDA reliably reproduces a known PAM requirement *in vitro* at adequate sequencing depth, we applied it to the full candidate panel. PAM signal calculated from NanoPAMDA was heterogeneous across the 14 candidates **(Fig. 3)** (**Supplementary Tables 11 and 14**). E10, a Cas9 effector from a mouse gut metagenome, produced a high-confidence experimental 5’-NGG-3’ logo consistent with its Protein2PAM prediction, whereas others showed diffuse or low-confidence signals without a clear consensus. E1, owing to its lower sequencing depth, displayed a patchy heatmap.

**Figure 2.**
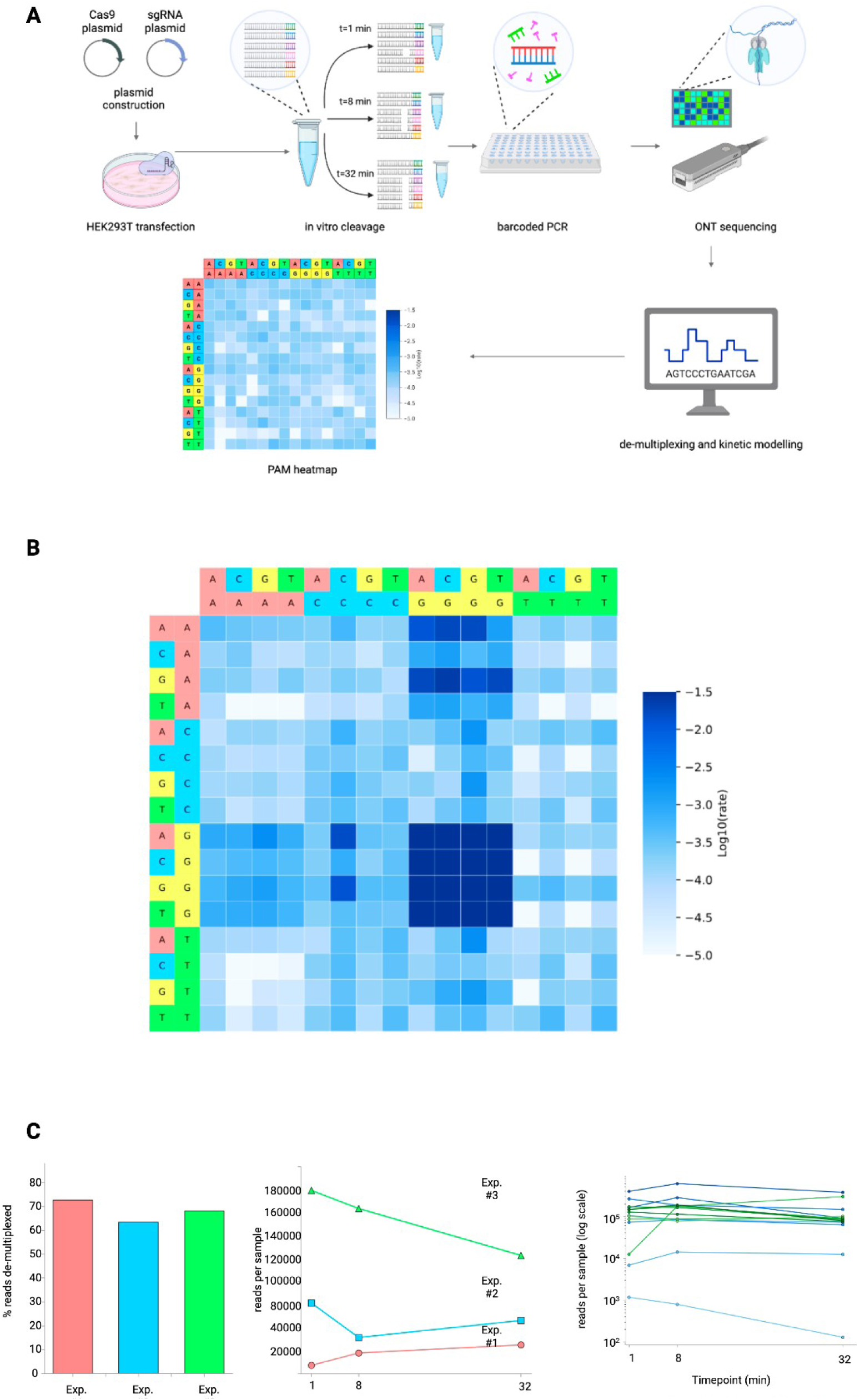
NanoPAMDA workflow, sequencing quality control, and PAM preference of SpCas9. **(A)** Overview of the NanoPAMDA experimental and computational workflow. Cas9 and sgRNA expression plasmids were co-transfected into HEK293T cells; cleared lysates were combined with a randomized PAM library in an in vitro cleavage reaction, with aliquots quenched at three timepoints (t = 1, 8, and 32 min). Cleavage products were amplified by barcoded PCR and sequenced on an Oxford Nanopore platform, then computationally demultiplexed and modeled kinetically to generate PAM preference heatmaps. **(B)** PAM preference heatmap for SpCas9, generated from NanoPAMDA cleavage kinetics. Log10-transformed pseudo-first-order cleavage rate constants are shown across the 4-nucleotide PAM window, plotted on a fixed color scale to enable direct comparison with other Cas9 variants. **(C)** Percentage of Nanopore reads successfully demultiplexed in each of three independent NanoPAMDA experiments, read depth per timepoint across the same three experiments, and read depth per timepoint for all 14 sequenced candidates across the three experiments (log scale), colored by candidate.

**Figure 3.**
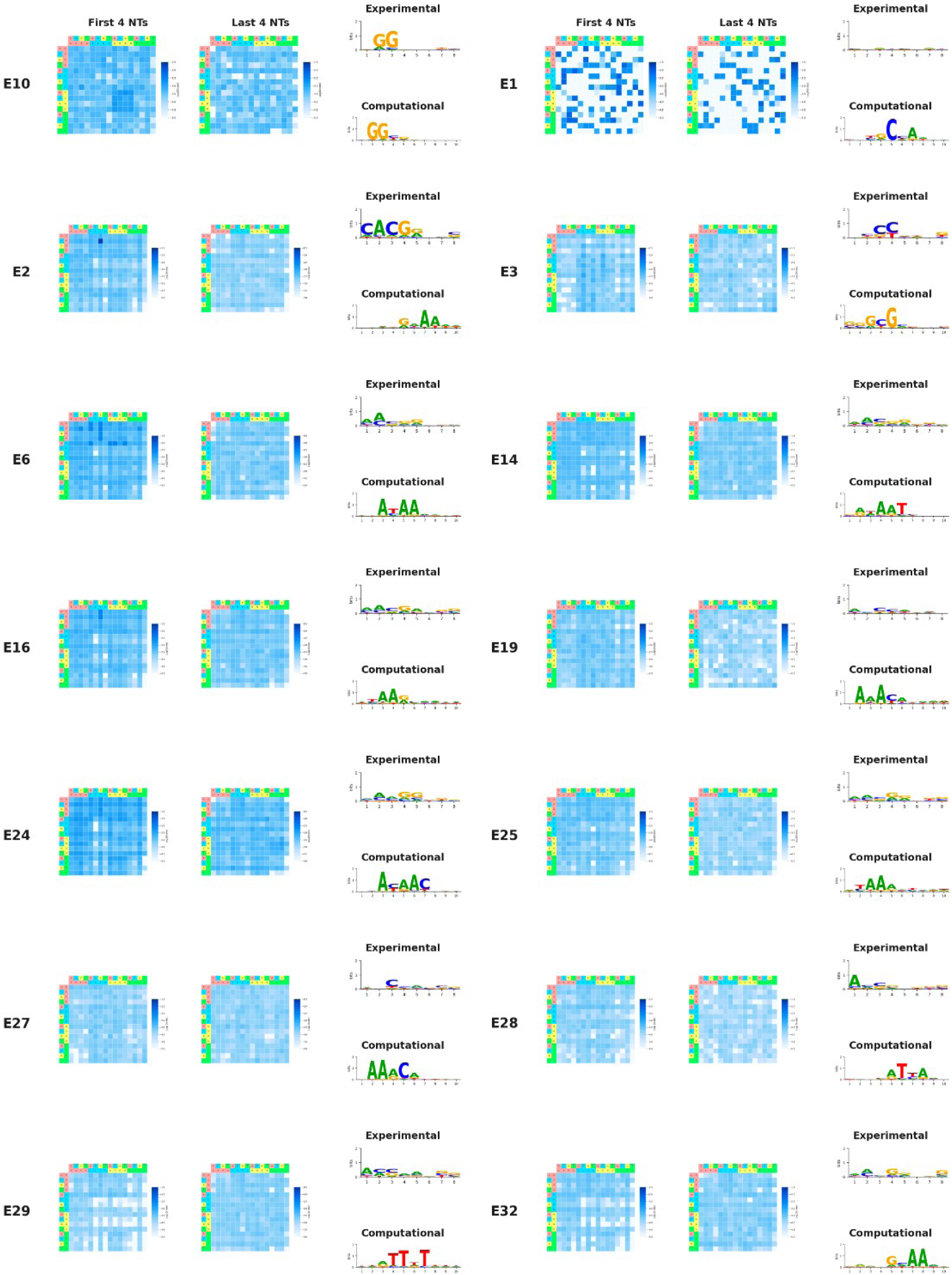
Experimental and predicted PAM preferences across the 14 Cas9 candidates. For each candidate (E10, E1, E2, E3, E6, E14, E16, E19, E24, E25, E27, E28, E29, E32), experimentally determined PAM preference heatmaps from NanoPAMDA cleavage kinetics are shown for the first and last 4 nucleotides of the PAM window (log10-transformed pseudo-first-order rate constants, blue scale; row and column headers colored by nucleotide identity as in Fig. 2B). For each candidate, an experimentally derived PAM sequence logo (Experimental, top) and the corresponding in silico Protein2PAM-predicted PAM logo (Computational, bottom) are shown side by side for direct comparison, with sequence logo height in bits.

### E10 is catalytically active in E. coli cells

Next, we tested whether E10 is catalytically active in bacteria when complexed with guides targeting genomic protospacers with its cognate 5’-NGG-3’ PAM. Because *E. coli* lacks an efficient non-homologous end-joining pathway, double-stranded breaks at a chromosomal target are generally lethal, so cleavage activity can be read out as a loss of colony formation. We therefore expressed E10 under an ATc-inducible promoter together with NGG sgRNAs targeting *adhE* (adhE-1, adhE-2), *pta* (pta-1, pta-2), or *xylB* genes, or a non-targeting (NT) sgRNA, and counted colonies with and without ATc **(Fig. 4) (Supplementary Table 15)**. The NT control formed comparable numbers of colonies in both conditions (2,920 ± 249 CFU without ATc; 3,840 ± 1,414 CFU with ATc), indicating that ATc-induced E10 expression is not by itself toxic in E. coli. In contrast, both *adhE*-targeting sgRNAs abolished colony formation upon induction (0 CFU in both replicates for each guide, versus 3,560 ± 668 and 3,884 ± 1,148 CFU without ATc), consistent with E10-mediated cleavage of the chromosomal target. The *pta*-targeting sgRNAs produced smaller and more variable reductions (pta-1, 4,256 ± 543 to 3,136 ± 1,595 CFU; pta-2, 1,972 ± 537 to 1,532 ± 979 CFU), which, with two replicates per condition, we cannot distinguish from the NT control. The *xylB* sgRNA yielded no colonies with or without ATc and was therefore uninformative. Together, these results suggest that E10 is catalytically active in *E. coli* under the tested conditions, with the clearest activity at *adhE*.

**Figure 4.**
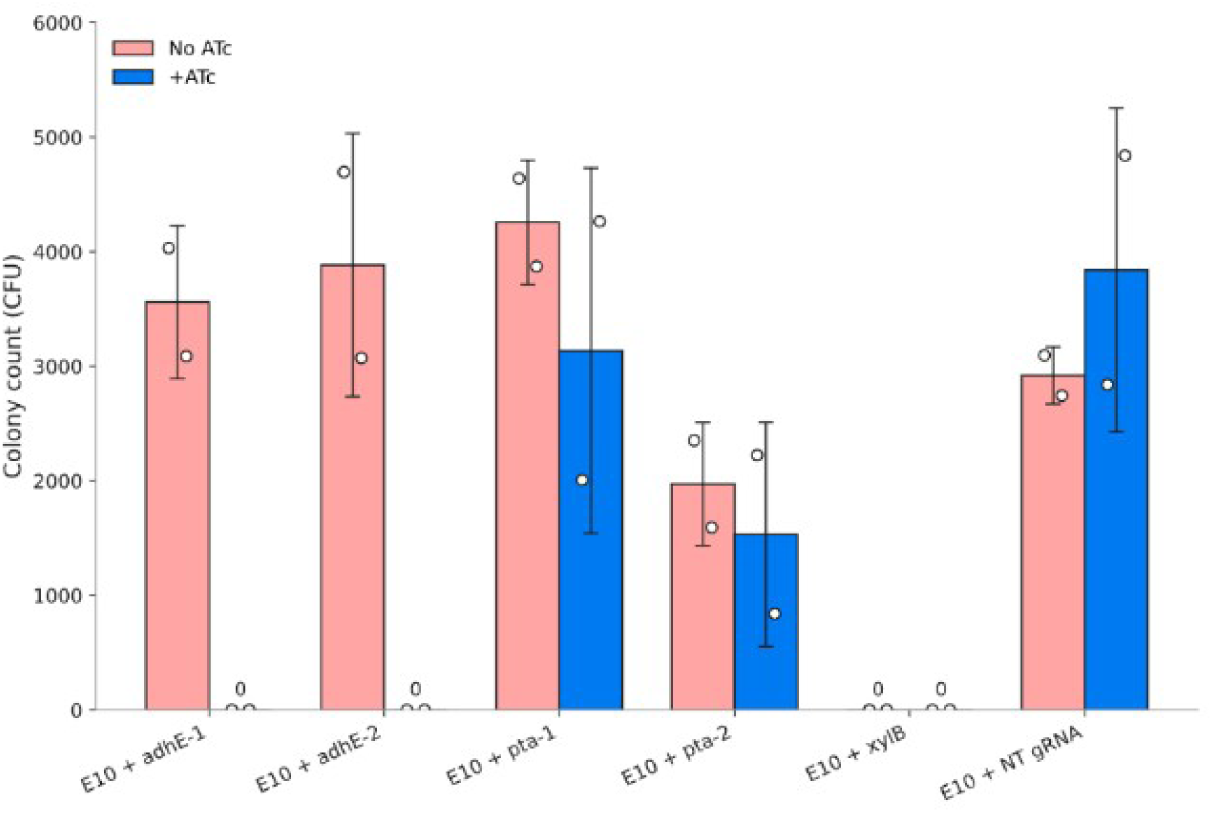
ATc-inducible E10 targeting of metabolic genes reduces colony formation in *E. coli*. Colony counts (CFU) of E. coli carrying E10 and a guide RNA targeting *adhE* (adhE-1, adhE-2), *pta* (pta-1, pta-2) or *xylB*, or a non-targeting (NT) gRNA control, plated without (pink) or with (blue) ATc to induce E10 expression. Bars show the mean and error bars the s.d. of n = 2 biological replicates; open circles show individual replicates. Zero colonies were recovered for both adhE guides with ATc and for xylB under both conditions.

## DISCUSSION

This study establishes an end-to-end pipeline for mining uncultivated metagenomic datasets for novel, structurally conserved CRISPR-Cas9 nucleases, and introduces NanoPAMDA, an Oxford Nanopore long-read adaptation of an *in vitro* PAM determination assay, for fast, scalable mapping of depletion kinetics across randomized target PAM libraries. Conventional HT-PAMDA relies on short-read Illumina sequencing, which imposes throughput bottlenecks, higher per-run costs, and longer turnaround times. NanoPAMDA’s single-step barcoded PCR coupled with Oxford Nanopore sequencing determined time-resolved depletion kinetics, with demultiplexing efficiencies of 63.4 to 72.7% and performance benchmarked directly against SpCas9 controls (**Fig. 2B–C**). This shows that NanoPAMDA and PCR-free long-read platforms can more broadly reach the depth needed for biochemical kinetic modeling in a fraction of the turnaround time of established methods.

In our hands, only 1 out of 14 (∼7%) metagenome-derived Cas9 effectors were functional in vitro and in E. coli. This rate is similar to the previously published ∼8.5% rate for Cas9 effectors derived from isolated single whole bacterial genomes, but significantly lower than the ∼70% success rate reported for metagenome-derived Cas9 effectors (Alexander et al., 2023; Butterfield et al., 2025). **The discrepancy** could be due to the relative abundance of human microbiome-derived Cas9 candidates in our datasets.

E10 is a mouse gut microbiome-derived nuclease (Xiao et al., 2015; PRJEB7759) sharing only about 36% amino acid identity with Streptococcus pyogenes Cas9 (SpCas9); its PAM-interacting domain shares approximately 23% amino acid identity with that of SpCas9. This level of divergence is typically too large to reliably infer PAM specificity from sequence alone, yet three independent lines of evidence converged on the same conclusion. Among six known Cas9 orthologs, the closest match of E10’s structure was SpCas9 (TM-score = 0.63; **Fig. 1E**). Protein2PAM independently predicted an NGG consensus at high confidence (0.904). NanoPAMDA confirmed this directly: E10’s five highest-ranked 4-nucleotide PAMs, AGGG, GGGA, TGGA, AGGT, and AGGC, were uniformly NGG-containing (**Fig. 3**). Taken together, these results show that E10 retains SpCas9-like PAM specificity despite its extensive sequence divergence, a conclusion made possible by direct NanoPAMDA measurement alongside the structural and computational evidence.

Several limitations of this study should be noted. Only one of the 14 candidates showed PAM-specific activity in vitro, so the panel provides limited power to compare success rates between data sources. The bacterial assay used two replicates per condition, depletion was clearest at a single locus (adhE), and the pta-targeting guides could not be distinguished from the non-targeting control; additional replicates and targets will be needed to quantify E10 activity. Finally, E10 activity in mammalian cells remains to be established.

We believe NanoPAMDA will enable researchers to characterize novel, functional CRISPR effector proteins using bespoke PAM library designs in a high-throughput, rapid manner.

## METHODS

### Metagenomic and Raw-Read Data Sources

Our computational pipeline was applied to a multi-biome collection of publicly available metagenome-assembled genomes (MAGs) and uncultivated viral contigs, aggregated from 17 metagenomic studies and resources spanning host-associated and environmental microbiomes, comprising 620,185 unique reconstructed genomes **(Supplementary Table 1)**.

Host-associated sources included the human gut microbiome **(Pasolli et al., 2019; Zeng et al., 2022; Forster et al., 2019; Almeida et al., 2021)**, human oral and vaginal microbiomes, and animal-associated gut catalogs spanning mouse **(Lesker et al., 2020)**, horse **(Li et al., 2023)**, pig, chicken, cow, honeybee, zebrafish, and non-model fish **(Gurbich et al., 2023)**, as well as a broad cross-species wild-animal gut resource **(Youngblut et al., 2020)**. Environmental sources included Earth’s microbiomes at large **(Nayfach et al., 2021)**, marine systems **(Xu et al., 2024; Nishimura & Yoshizawa, 2022)**, freshwater systems **(Garner et al., 2023; Borton et al., 2025)**, soil **(Ma et al., 2023)**, the Tibetan Plateau aquatic microbiome **(Cheng et al., 2024)**, thawing permafrost **(Woodcroft et al., 2018)**, and activated sludge from wastewater treatment **(Ye et al., 2020)**.

Raw shotgun metagenomic sequencing reads were obtained from two human microbiome studies deposited in the database of Genotypes and Phenotypes (dbGaP): the NIH Human Microbiome Project Core Microbiome Sampling Protocol A (HMP-A; dbGaP accession phs000228.v4.p1) and the Trans-omics for Precision Dentistry and Early Childhood Caries study (TOPDECC; Genome-Wide Genotyping and Microbiome in the ZOE 2.0 Study; dbGaP accession phs002232.v1.p1) **(Supplementary Table 2)**. Raw reads from both studies were assembled de novo using MetaHipMer2 (Hofmeyr et al., 2020).

### CRISPR-Cas9 Candidate Discovery

Assemblies were searched for CRISPR arrays using PILER-CR (v1.06) (Edgar, 2007). Open reading frames (ORFs) and protein translations were predicted using Prodigal (v2.6.3) in metagenomic mode (-p meta) **(Hyatt et al., 2010)**. The predicted protein sequences were queried against a curated Cas profile Hidden Markov Models **(Makarova et al., 2015)** using the hmmsearch module from the HMMER (v3.3.2) suite **(Eddy, 2011)**. A putative Cas protein sequence was retained only if it met all of the following alignment thresholds: alignment coverage >70%, bitscore >= 40, e-value < 1e-5.

Candidate tracrRNAs associated with each Cas9 were identified by extracting the 20 kb flanking region surrounding each candidate locus and analyzing these sequences using CRISPRtracrRNA with default parameters to predict candidate tracrRNAs for each locus **(Mitrofanov et al., 2022)**. Candidate sgRNA architectures were retained only if the tracrRNA fell within the 70–150 nucleotide length window. The resulting loci are listed in Supplementary Tables 3 and 5.

Environmental candidates were clustered using CD-HIT (Li & Godzik, 2006) at a sequence identity threshold of 50% (-c 0.50 -n 3); human microbiome candidates were clustered at a sequence identity threshold of 90% (-c 0.90 -n 5). Cluster representatives are in Supplementary Tables 4 and 6. For visualization purposes **(Fig. 1C)**, the combined length-filtered candidate pool was independently re-clustered at 90% sequence identity using CD-HIT **(Li & Godzik, 2006)**. The representatives were combined with the 14 candidates selected for experimental characterization and six known Cas9 orthologs (SpCas9, FnCas9, Nme2Cas9, Cas9d, SauCas9, and CjCas9; 1,125 total sequences), aligned using MAFFT **(Katoh & Standley, 2013)** (default parameters), and used to construct a phylogenetic tree with FastTree **(Price et al., 2009). Environmental cluster representatives** were screened for sequence novelty via the NCBI web BLAST server (blastp, non-redundant protein database) **(Boratyn et al., 2013)**. Conservation of critical catalytic residues was assessed by aligning each candidate against a panel of six known Cas9 orthologs (SpCas9, FnCas9, Nme2Cas9, Cas9d, SauCas9, and CjCas9) using Clustal Omega (EMBL-EBI web server) **(Sievers et al., 2011)**. Within this multiple sequence alignment, columns corresponding to functionally critical RuvC, HNH, and bridge-helix/PI catalytic residues were identified using SpCas9 residues: D10; S15, N15, or Y15; R66; R70 or K70; R74, T74, or D74; R78 or L78; R165 or E165; E762 or D762; H840; N854; N863; H982 or S982; H983; and D986. Global structural similarity to characterized Cas9 orthologs was assessed by predicting each candidate’s three-dimensional structure using ESMFold (**Lin et al., 2023**) and computing pairwise structural alignments against reference Cas9 structures using the Foldseek (**van Kempen et al., 2024**) (**Supplementary Figure 2; Supplementary Table 9**).

Two-dimensional secondary structures of the tracrRNA tails of the surviving subset were predicted using the NUPACK web server (default parameters) (**Zadeh et al., 2011**), and candidates were retained only if the structure showed a stable hairpin, proper nexus architecture, and a distinct nexus junction. Structures for the 7 environmental microbiome-derived candidates are provided in **Supplementary Figure 1**.

Human microbiome cluster representatives were evaluated for conservation of critical catalytic residues as described above; the 2D guide RNA structural rules and structural similarity criteria were not applied to this arm. Candidates were evaluated sequentially against the catalytic residue criterion until 7 candidates satisfying the filter were identified **(Supplementary Tables 6 and 8).**

### In Silico PAM Prediction

In silico predictions were generated for the 14 candidates **(Supplementary Table 11)** selected for experimental testing using Protein2PAM (Nayfach et al., 2026). To assess whether the 14 candidates were representative of the broader PAM-prediction landscape, Protein2PAM predictions were extended to all 491 candidate representatives from the selection-stage clustering (424 environmental, 50% identity clusters; 67 human microbiome, 90% identity clusters) (Supplementary Table 10).

### Molecular Cloning

The RTW3027 plasmid (Addgene #139987) was linearized by AgeI and NotI (New England Biolabs). The complete, codon-optimized sequence of each Cas9 ortholog was synthesized as three overlapping fragments (gBlocks, Integrated DNA Technologies) **(Supplementary Tables 7, 8 and 12)**. A separate P2A-EGFP reporter cassette was generated via PCR amplification. The final mammalian expression vector was constructed via multi-fragment isothermal assembly (Gibson Assembly). The BPK1520 gRNA scaffold vector (Addgene #65777) was first digested using HindIII and BsmBI, followed by the insertion of a synthesized scaffold gBlock fragment. The resulting vector was digested with BsmBI, and the HT-PAMDA library 1 spacer was cloned into it.

### Cell Culture

Human embryonic kidney (HEK293T) cells (ATCC CRL-11268) were cultured and maintained in High-Glucose Dulbecco’s Modified Eagle Medium (DMEM; Gibco) supplemented with 10% fetal bovine serum (FBS; Gibco) and 1% penicillin-streptomycin (10,000 U/mL; Gibco). Cells were grown in a humidified incubator maintained at 37°C with 5% CO2. Cultures were monitored daily and routinely passaged upon reaching 80–90% confluence using 0.25% Trypsin-EDTA (Gibco) for cell dissociation.

### Transfection

For functional in vivo expression and endogenous ribonucleoprotein (RNP) assembly, 1.5 × 10⁵ HEK293T cells were seeded into 24-well tissue culture plates and allowed to reach 70–80% confluence at the time of transfection. Transient co-transfections were performed in a total volume of 30 μL per well using Opti-MEM Reduced Serum Medium (Thermo Fisher Scientific) prepared with the following components: 500 ng effector plasmid, 165 ng gRNA plasmid, and 4 μL of CanFast Transfection Reagent (Canvax, Spain). The transfected cells were incubated under standard growth conditions (37°C, 5% CO2) without a post-transfection media change. Cells were harvested 48 hours post-transfection for downstream lysis.

### In vitro Cleavage Assay and Nanopore Sequencing

Cleavage assays were performed as previously described (Walton et al., 2021). The remaining uncleaved, intact plasmid library fractions from each quenched time point were amplified in a single reaction using custom PCR primers (**Supplementary Table 13**). The library (Addgene #160132) was purified and prepared for sequencing using the Native Barcoding Kit 96 V14 (SQK-NBD114.96; Oxford Nanopore Technologies). A single native ONT barcode was applied to the entire pooled reaction. The finalized library was loaded onto a MinION flow cell (Flo-MIN114; R10.4.1 chemistry, Oxford Nanopore Technologies) and sequenced on a MinION Mk1D instrument. POD5 files were processed with the Dorado basecaller (v1.4.0, Oxford Nanopore Technologies), using the super-accurate (sup) model. NanoPAMDA pipeline binned reads based on their unique 10-nucleotide forward and reverse index pairs. A maximum threshold of 2 mismatches (≤ 2 errors) was permitted for a successful barcode alignment. Reads successfully mapping to a paired index combination within the error threshold were extracted and grouped into distinct sample-specific FASTQ files. NanoPAMDA searched for an exact match to the specific 20-nucleotide target spacer sequence (e.g., SPACER1: GGGCACGGGCAGCTTGCCGG) and extracted the immediately adjacent 8-nucleotide sequence at the 3’ end, which represents the putative PAM. Reads containing ambiguous base calls (N) within this extracted window were discarded.

Nanopore cleavage timepoint data were normalized against a high-depth untreated control library. Because kinetic fitting requires a defined core PAM window, the 8-nucleotide window was split into two 4-nucleotide marginal windows, which were normalized, fit, and visualized independently. A pseudo-first-order kinetic model was applied to the normalized sequence counts. The depletion of each specific PAM motif over time was fitted to a standard exponential decay function: y(t) = a × e^(−kt), where y(t) represents the normalized read fraction of the PAM at time t, a is the initial read fraction at t=0, k is the pseudo-first-order rate constant of cleavage, and t is the reaction time converted to seconds (0, 60, 480, and 1920 seconds). The optimal rate constant (k) for each sequence was determined using a non-linear least-squares curve-fitting algorithm (SciPy) **(Virtanen et al., 2020)**. Sequences that failed to meet a strict minimum threshold of 4 total reads across the dataset, or that had missing valid counts at least one time point, were excluded from the regression analysis and assigned a null value. The calculated cleavage rate constants (k) were averaged across replicates and spacer targets, log-transformed, and plotted on a standard quantitative color matrix. Heatmaps were constrained to a fixed scale (ranging from −5.0 to −1.5) to enable direct visual comparison of nuclease affinities across different experimental libraries and Cas9 variants.

### Bacterial Nuclease Activity Assay

The E10 Cas9 nuclease coding sequence was cloned into the pdCas9-bacteria backbone (Addgene #44249), replacing dCas9, using Gibson assembly. An E10-compatible gRNA scaffold was introduced into psgRNA (Addgene #114005), and sgRNAs targeting two sites each in *adhE* and *pta*, a site in *xylB*, and a non-targeting control were cloned using BsmBI-mediated ligation (**Supplementary Table 15**). Chemically competent *E. coli* Stbl3 cells were co-transformed with 80 ng E10 plasmid and 20 ng sgRNA plasmid, recovered in SOC medium, and plated on selective agar containing chloramphenicol (25 µg/mL) and kanamycin (50 µg/mL), with or without ATc (0.1 ng/mL) to induce E10 expression. Plates were incubated for 16 hours at 37 °C, and colonies were counted for two replicates per condition.

## Code and Data Availability

All custom scripts used in this study, including data ingestion and CRISPR array/Cas9 discovery scripts, guide RNA resolution and clustering scripts, novelty and structural validation scripts, PAM prediction protocols, and the full HT-PAMDA/Nanopore analysis pipeline, are available at https://github.com/igenel/NanoPAMDA/. The 17 public metagenome-assembled genome (MAG) catalogs used in this study, along with their citations and genome counts, are listed in Supplementary Table 1. Processed Cas9 candidate metadata tables (tracrRNA-length-filtered, clustered, and final selected candidate sets for both the environmental and human microbiome arms) are provided as Supplementary Data. Demultiplexed Nanopore sequencing data FASTQ generated for HT-PAMDA characterization in this study have been deposited in the Sequence Read Archive (SRA) under accession **PRJNA1504502**.

The human microbiome datasets analyzed in this study were obtained through authorized access to the NIH database of Genotypes and Phenotypes (dbGaP), under Project #34434. The datasets included the Human Microbiome Project Core Microbiome Sampling Protocol A (HMP-A; accession phs000228.v4.p1) and the Trans-omics for Precision Dentistry and Early Childhood Caries (TOPDECC/ZOE 2.0; accession phs002232.v1.p1). The original study data are available through dbGaP, with access to controlled data subject to approval by the relevant Data Access Committees and applicable data-use restrictions.

## Acknowledgement

The authors thank Osman Nuri Polat, Ömer Faruk Ağca, and CureCraft for their financial and research support.

We acknowledge the investigators and institutions responsible for generating and sharing the human microbiome datasets through the NIH database of Genotypes and Phenotypes (dbGaP).

The NIH Human Microbiome Project (HMP-A) was supported by the NIH Roadmap for Medical Research. **(Human Microbiome Project Consortium, 2012)** We acknowledge the contributions of Baylor College of Medicine and Washington University School of Medicine to the clinical data, as well as the Baylor College of Medicine Human Genome Sequencing Center, the Broad Institute, the Genome Center at Washington University, and the J. Craig Venter Institute to sequencing. We also acknowledge EMMES Corporation for its role in clinical data coordination and submission.

The ZOE 2.0 study, part of TOPDECC, was supported by the National Institute of Dental and Craniofacial Research (U01-DE025046). Genotyping was performed by the Center for Inherited Disease Research (CIDR), supported by NIDCR grant X01-HG010871.

## Contributions

Wet laboratory experiments were performed by I.G., except for the functional characterization in bacteria, which was performed by C.S. Computational analyses were performed by I.G. The study was conceived and designed by I.C.K. and I.G. I.C.K. supervised the work. I.C.K. and I.G. wrote the initial draft of the manuscript, and all authors contributed to reviewing and editing the manuscript.

## REFERENCES

Alexander, L.M., Aliaga Goltsman, D.S., Liu, J., Lin, J.-L., Temoche-Diaz, M.M., Laperriere, S.M., Neerincx, A., Bednarski, C., Knyphausen, P., Cohnen, A., Albers, J., Gonzalez-Osorio, L., Fregoso Ocampo, R., Oki, J., Devoto, A.E., Castelle, C.J., Lamothe, R.C., Cost, G.J., Butterfield, C.N., Thomas, B.C., Brown, C.T. (2023). Novel and engineered type II CRISPR systems from uncultivated microbes with broad genome editing capability. CRISPR J. 6, 261–277.

Almeida, A., Nayfach, S., Boland, M., et al. (2021). A unified catalog of 204,938 reference genomes from the human gut microbiome. Nat. Biotechnol. 39, 105–114.

Altae-Tran, H., Kannan, S., Demircioglu, F.E., Oshiro, R., Nety, S.P., McKay, L.J., Dlakić, M., Inskeep, W.P., Makarova, K.S., Macrae, R.K., Koonin, E.V., Zhang, F. (2021). The widespread IS200/IS605 transposon family encodes diverse programmable RNA-guided endonucleases. Science 374, 57–65.

Boratyn, G.M. et al. (2013). BLAST: a more efficient report with usability improvements. Nucleic Acids Res. 41, W29–W33.

Borton, M.A. et al. (2025). A functional microbiome catalog crowdsourced from North American rivers. Nature.

Butterfield, G.L., Rohm, D., Roberts, A., Nethery, M.A., Rizzo, A.J., Morone, D.J., Garnier, L., Iglesias, N., Barrangou, R., Gersbach, C.A. (2025). Characterization of diverse Cas9 orthologs for genome and epigenome editing. Proc. Natl. Acad. Sci. U.S.A. 122, e2417674122.

Cheng, M. et al. (2024). A genome and gene catalog of the aquatic microbiomes of the Tibetan Plateau. Nat. Commun.

Cong, L., Ran, F.A., Cox, D., Lin, S., Barretto, R., Habib, N., Hsu, P.D., Wu, X., Jiang, W., Marraffini, L.A., Zhang, F. (2013). Multiplex genome engineering using CRISPR/Cas systems. Science 339, 819–823.

Doudna, J.A., Charpentier, E. (2014). Genome editing. The new frontier of genome engineering with CRISPR-Cas9. Science 346, 1258096.

Eddy, S.R. (2011). Accelerated profile HMM searches. PLoS Comput. Biol. 7, e1002195.

Edgar, R.C. (2007). PILER-CR: fast and accurate identification of CRISPR repeats. BMC Bioinformatics 8, 18.

Edraki, A., Mir, A., Ibraheim, R., Gainetdinov, I., Yoon, Y., Song, C.Q., Cao, Y., Gallant, J., Xue, W., Rivera-Pérez, J.A., et al. (2019). A compact, high-accuracy Cas9 with a dinucleotide PAM for in vivo genome editing. Mol. Cell 73, 714–726.e4.

Fang, T., Bogensperger, L., Feer, L., Allam, A., Balázs, Z., von Mering, C., Sunagawa, S., Krauthammer, M., Schwank, G. (2026). Uncovering Cas9 PAM diversity through metagenomic mining and machine learning. Nat. Commun. 17, 2510.

Forster, S.C., Kumar, N., Anonye, B.O., et al. (2019). A human gut bacterial genome and culture collection for improved metagenomic analyses. Nat. Biotechnol. 37, 186–192.

Garner, R.E. et al. (2023). A genome catalogue of lake bacterial diversity and its drivers at continental scale. Nat. Microbiol.

Gasiunas, G., Young, J.K., Karvelis, T., Kazlauskas, D., Urbaitis, T., Jasnauskaite, M., Grusyte, M.M., Paulraj, S., Wang, P.H., Hou, Z., Dooley, S.K., Cigan, M., Alarcon, C., Chilcoat, N.D., Bigelyte, G., Curcuru, J.L., Mabuchi, M., Sun, Z., Fuchs, R.T., Schildkraut, E., Weigele, P.R., Jack, W.E., Robb, G.B., Venclovas, C., Siksnys, V. (2020). A catalogue of biochemically diverse CRISPR-Cas9 orthologs. Nat. Commun. 11, 5512.

Gurbich, T.A. et al. (2023). MGnify Genomes: a resource for biome-specific microbial genome catalogues. J. Mol. Biol.

Hofmeyr, S., Egan, R., Georganas, E., et al. (2020). Terabase-scale metagenome coassembly with MetaHipMer. Sci. Rep. 10, 10689.

Hsiau, T., Conant, D., Rossi, N., Maures, T., Waite, K., Yang, J., Joshi, S., Kelso, R., Holden, K., Enzmann, B.L., Stoner, R. (2019). Inference of CRISPR Edits from Sanger trace data. bioRxiv. 10.1101/251082.

Human Microbiome Project Consortium (2012). Structure, function and diversity of the healthy human microbiome. Nature 486, 207–214

Hyatt, D., Chen, G.L., LoCascio, P.F., Land, M.L., Larimer, F.W., Hauser, L.J. (2010). Prodigal: prokaryotic gene recognition and translation initiation site identification. BMC Bioinformatics 11, 119.

Jinek, M., Chylinski, K., Fonfara, I., Hauer, M., Doudna, J.A., Charpentier, E. (2012). A programmable dual-RNA-guided DNA endonuclease in adaptive bacterial immunity. Science 337, 816–821.

Katoh, K., Standley, D.M. (2013). MAFFT multiple sequence alignment software version 7: improvements in performance and usability. Mol. Biol. Evol. 30, 772–780.

Kim, E., Koo, T., Park, S.W., Kim, D., Kim, K., Cho, H.Y., Song, D.W., Lee, K.J., Jung, M.H., Kim, S., Kim, J.H., Kim, J.H., Kim, J.S. (2017). In vivo genome editing with a small Cas9 orthologue derived from *Campylobacter jejuni*. Nat. Commun. 8, 14500.

Lesker, T.R. et al. (2020). An integrated metagenome catalog reveals new insights into the murine gut microbiome. Cell Rep.

Li, W., Godzik, A. (2006). Cd-hit: a fast program for clustering and comparing large sets of protein or nucleotide sequences. Bioinformatics 22, 1658–1659.

Li, F. et al. (2023). Expanded catalogue of metagenome-assembled genomes reveals resistome characteristics and athletic performance-associated microbes in horse. Microbiome.

Lin, Z., Akin, H., Rao, R., Hie, B., Zhu, Z., Lu, W., Smetanin, N., Verkuil, R., Kabeli, O., Shmueli, Y., dos Santos Costa, A., Fazel-Zarandi, M., Sercu, T., Candido, S., Rives, A. (2023). Evolutionary-scale prediction of atomic-level protein structure with a language model. Science 379, 1123–1130.

Ma, B. et al. (2023). A genomic catalogue of soil microbiomes boosts mining of biodiversity and genetic resources. Nat. Commun.

Makarova, K.S. et al. (2015). An updated evolutionary classification of CRISPR–Cas systems. Nat. Rev. Microbiol. 13, 722–736.

Mali, P., Yang, L., Esvelt, K.M., Aach, J., Guell, M., DiCarlo, J.E., Norville, J.E., Church, G.M. (2013). RNA-guided human genome engineering via Cas9. Science 339, 823–826.

Mitrofanov, A., Ziemann, M., Alkhnbashi, O.S., Hess, W.R., Backofen, R. (2022). CRISPRtracrRNA: robust approach for CRISPR tracrRNA detection. Bioinformatics 38(Suppl_2), ii42–ii48.

Nayfach, S., Roux, S., Seshadri, R., et al. (2021). A genomic catalog of Earth’s microbiomes. Nat. Biotechnol. 39, 499–509.

Nayfach, S. et al. (2026). Customizing CRISPR–Cas PAM specificity with protein language models. Nat. Biotechnol.

Nishimasu, H., Ran, F.A., Hsu, P.D., Konermann, S., Shehata, S.I., Dohmae, N., Ishitani, R., Zhang, F., Nureki, O. (2014). Crystal structure of Cas9 in complex with guide RNA and target DNA. Cell 156, 935–949.

Nishimura, Y., Yoshizawa, S. (2022). The OceanDNA MAG catalog contains over 50,000 prokaryotic genomes originated from various marine environments. Sci. Data.

Pasolli, E., Asnicar, F., Manara, S., et al. (2019). Extensive unexplored human microbiome diversity revealed by over 150,000 genomes from metagenomes spanning age, geography, and lifestyle. Cell 176, 649–662.

Price, M.N., Dehal, P.S., Arkin, A.P. (2009). FastTree: computing large minimum evolution trees with profiles instead of a distance matrix. Mol. Biol. Evol. 26, 1641–1650.

Ran, F.A., Cong, L., Yan, W.X., Scott, D.A., Gootenberg, J.S., Kriz, A.J., Zetsche, B., Shalem, O., Wu, X., Makarova, K.S., Koonin, E.V., Sharp, P.A., Zhang, F. (2015). In vivo genome editing using *Staphylococcus aureus* Cas9. Nature 520, 186–191.

Xiao, L., Feng, Q., Liang, S., Sonne, S.B., Xia, Z., Qiu, X., Li, X., Long, H., Zhang, J., Zhang, D., Liu, C., Fang, Z., Chou, J., Glanville, J., Hao, Q., Kotowska, D., Colding, C., Licht, T.R., Wu, D., Yu, J., Sung, J.J.Y., Liang, Q., Li, J., Jia, H., Lan, Z., Tremaroli, V., Dworzynski, P., Nielsen, H.B., Bäckhed, F., Doré, J., Le Chatelier, E., Ehrlich, S.D., Lin, J.C., Arumugam, M., Wang, J., Madsen, L., Kristiansen, K. (2015). A catalog of the mouse gut metagenome. Nat. Biotechnol. 33, 1103–1108.

Sievers, F., Wilm, A., Dineen, D., Gibson, T.J., Karplus, K., Li, W., Lopez, R., McWilliam, H., Remmert, M., Söding, J., Thompson, J.D., Higgins, D.G. (2011). Fast, scalable generation of high-quality protein multiple sequence alignments using Clustal Omega. Mol. Syst. Biol. 7, 539.

Xu, S., Huang, H., Chen, S., Muhammad, Z.U.A., Wei, W., Xie, W., Jiang, H., Hou, S. (2024). Recovery of 1887 metagenome-assembled genomes from the South China Sea. Sci. Data 11, 197.

van Kempen, M., Kim, S.S., Tumescheit, C., Mirdita, M., Lee, J., Gilchrist, C.L.M., Söding, J., Steinegger, M. (2024). Fast and accurate protein structure search with Foldseek. Nat. Biotechnol. 42, 243–246.

Virtanen, P., Gommers, R., Oliphant, T.E., et al. (2020). SciPy 1.0: fundamental algorithms for scientific computing in Python. Nat. Methods 17, 261–272.

Walton, R.T., Christie, K.A., Whittaker, M.N., Kleinstiver, B.P. (2020). Unconstrained genome targeting with near-PAMless engineered CRISPR-Cas9 variants. Science 368, 290–296.

Walton, R.T., Hsu, J.Y., Joung, J.K., Kleinstiver, B.P. (2021). Scalable characterization of the PAM requirements of CRISPR-Cas enzymes using HT-PAMDA. Nat. Protoc. 16, 1511–1547.

Woodcroft, B.J. et al. (2018). Genome-centric view of carbon processing in thawing permafrost. Nature.

Ye, L. et al. (2020). Machine learning-aided analyses of thousands of draft genomes reveal specific features of activated sludge processes. Microbiome.

Youngblut, N.D. et al. (2020). Large-scale metagenome assembly reveals novel animal-associated microbial genomes, biosynthetic gene clusters, and other genetic diversity. mSystems.

Zadeh, J.N., Steenberg, C.D., Bois, J.S., Wolfe, B.R., Pierce, M.B., Khan, A.R., Dirks, R.M., Pierce, N.A. (2011). NUPACK: analysis and design of nucleic acid systems. J. Comput. Chem. 32, 170–173.

Zeng, S. et al. (2022). A compendium of 32,277 metagenome-assembled genomes and over 80 million genes from the early-life human gut microbiome. Nat. Commun.

