## Supplementary Figure1 for "NanoPAMDA enables rapid PAM characterization of metagenome-derived Cas9 effectors"

### 3300010270-21_contig-100_98

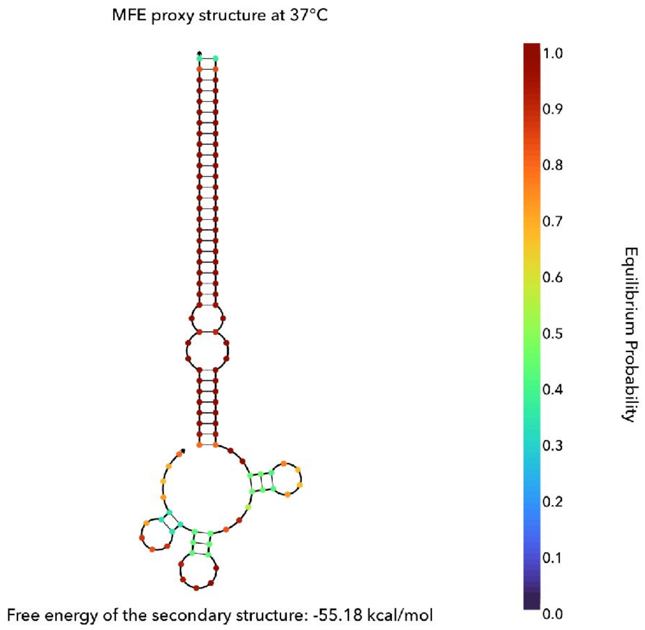


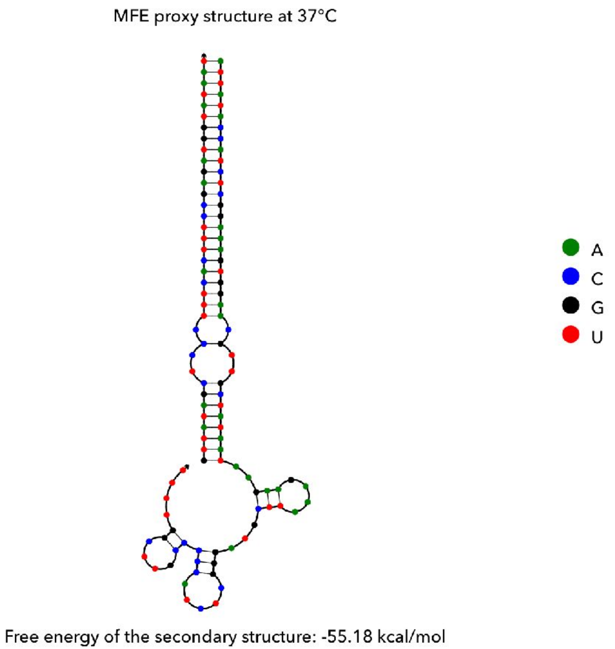

### 3300017518-52_contig-100_25

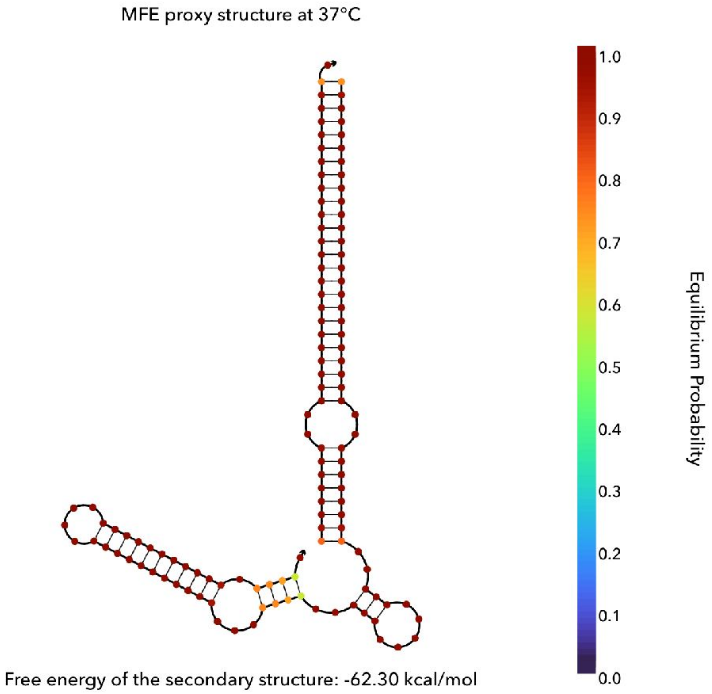


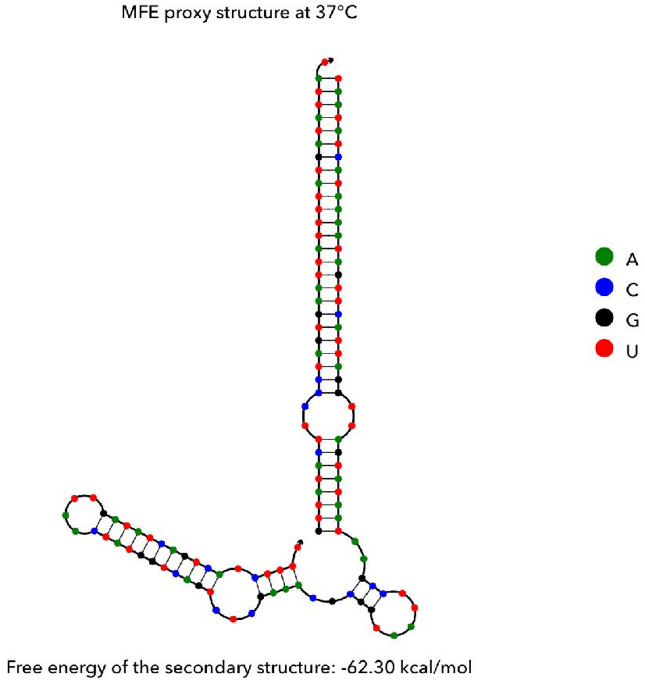

### 3300025572-13_contig-100_171

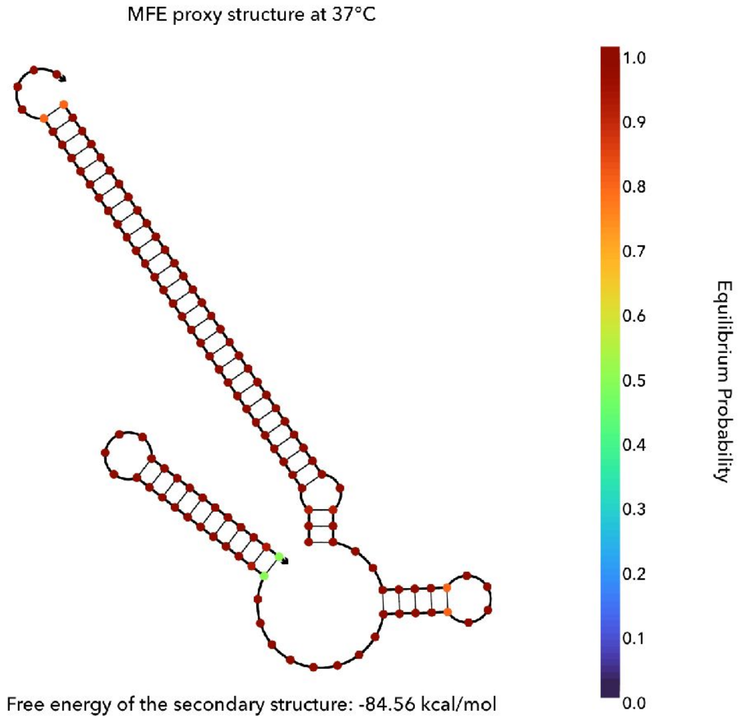


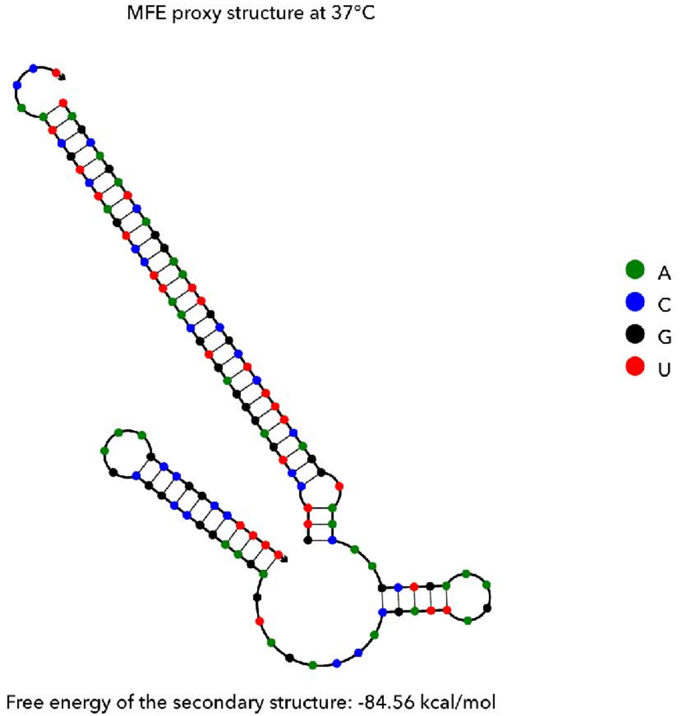

### MGYG000002579_contig-100_7

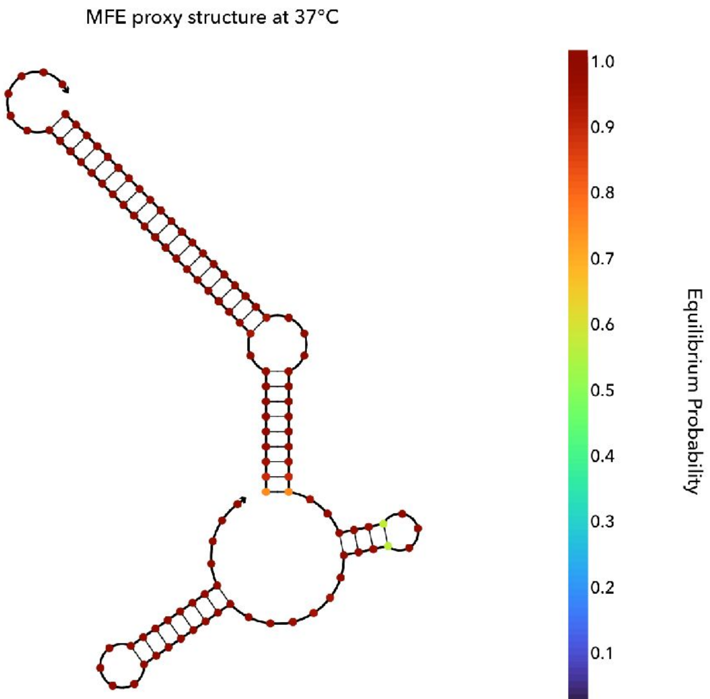


Free energy of the secondary structure: -57.78 kcal/mol

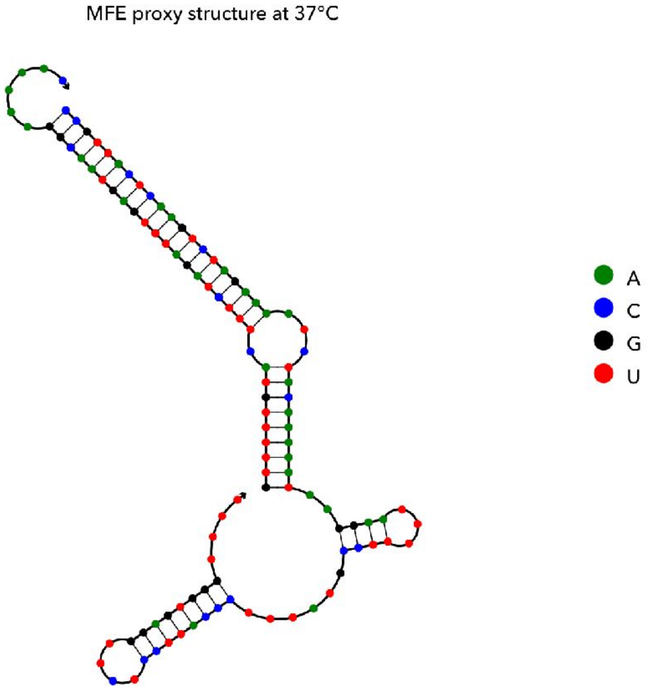


Free energy of the secondary structure: -57.78 kcal/mol

### XieH-2016--YSZC12003-36058--bin-50_contig-100_43

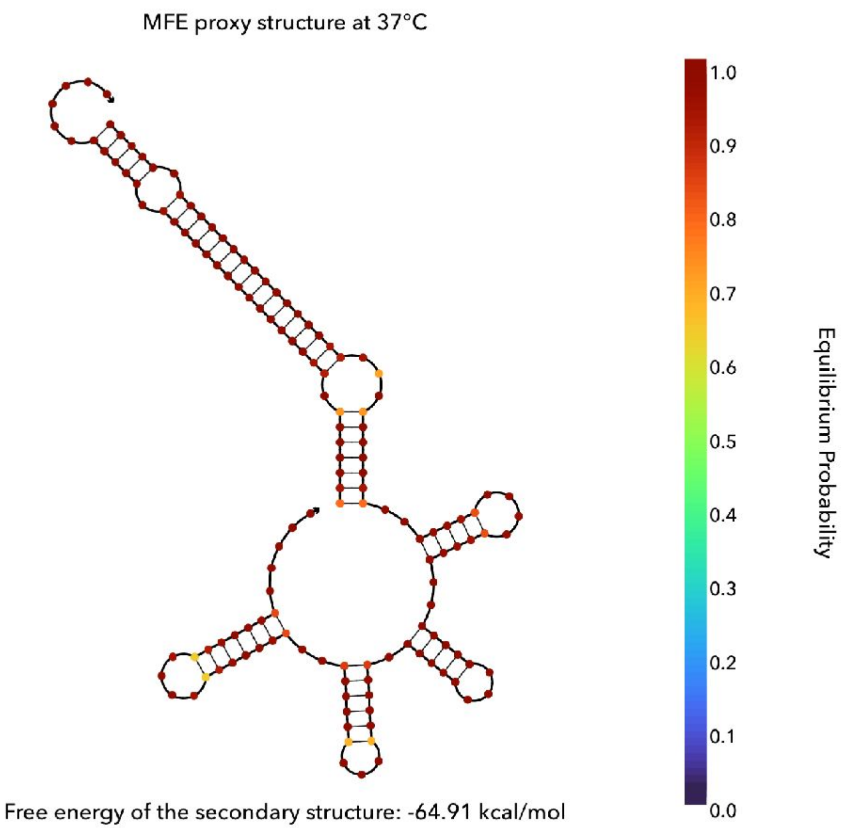


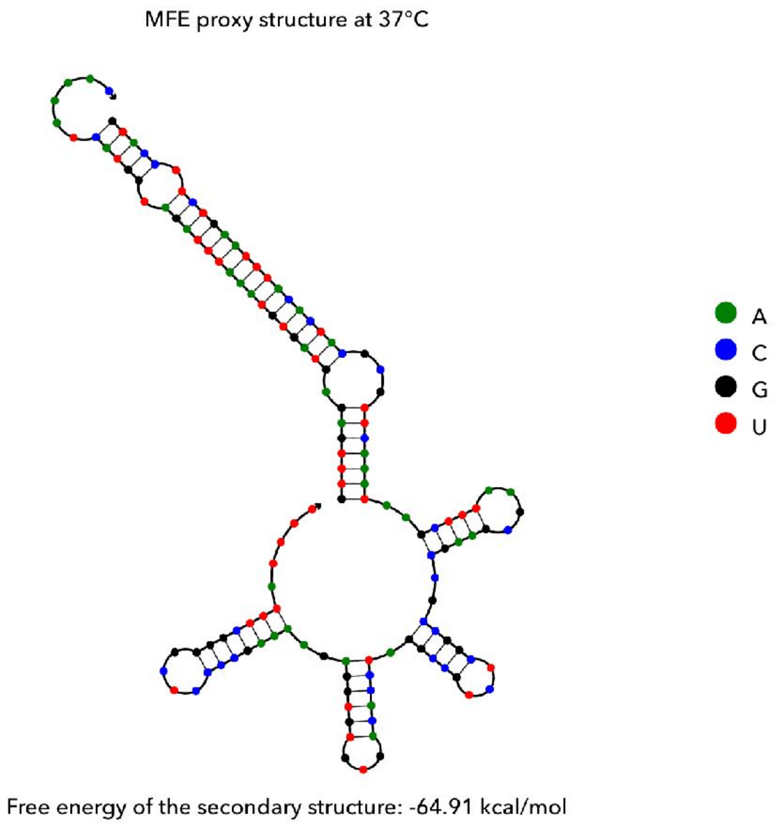

### ZeeviD-2015--PNP-Main-540--bin-18_contig-100_24

MFE proxy structure at 37°C

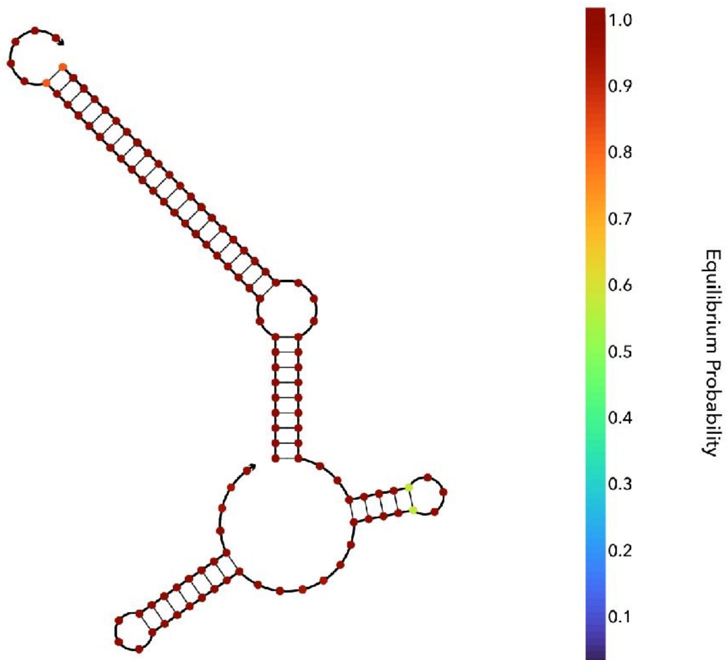


Free energy of the secondary structure: -60.57 kcal/mol

MFE proxy structure at 37°C

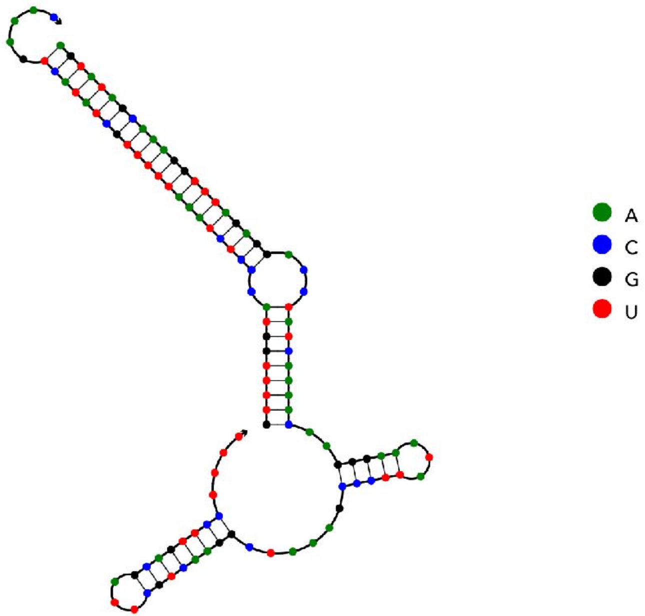


Free energy of the secondary structure: -60.57 kcal/mol
