## Supplementary figures and images for "NanoPAMDA enables rapid PAM characterization of metagenome-derived Cas9 effectors"

### Supplementary Figure2

#### ESMFold-predicted structures of the 6 MAG-derived validated candidates

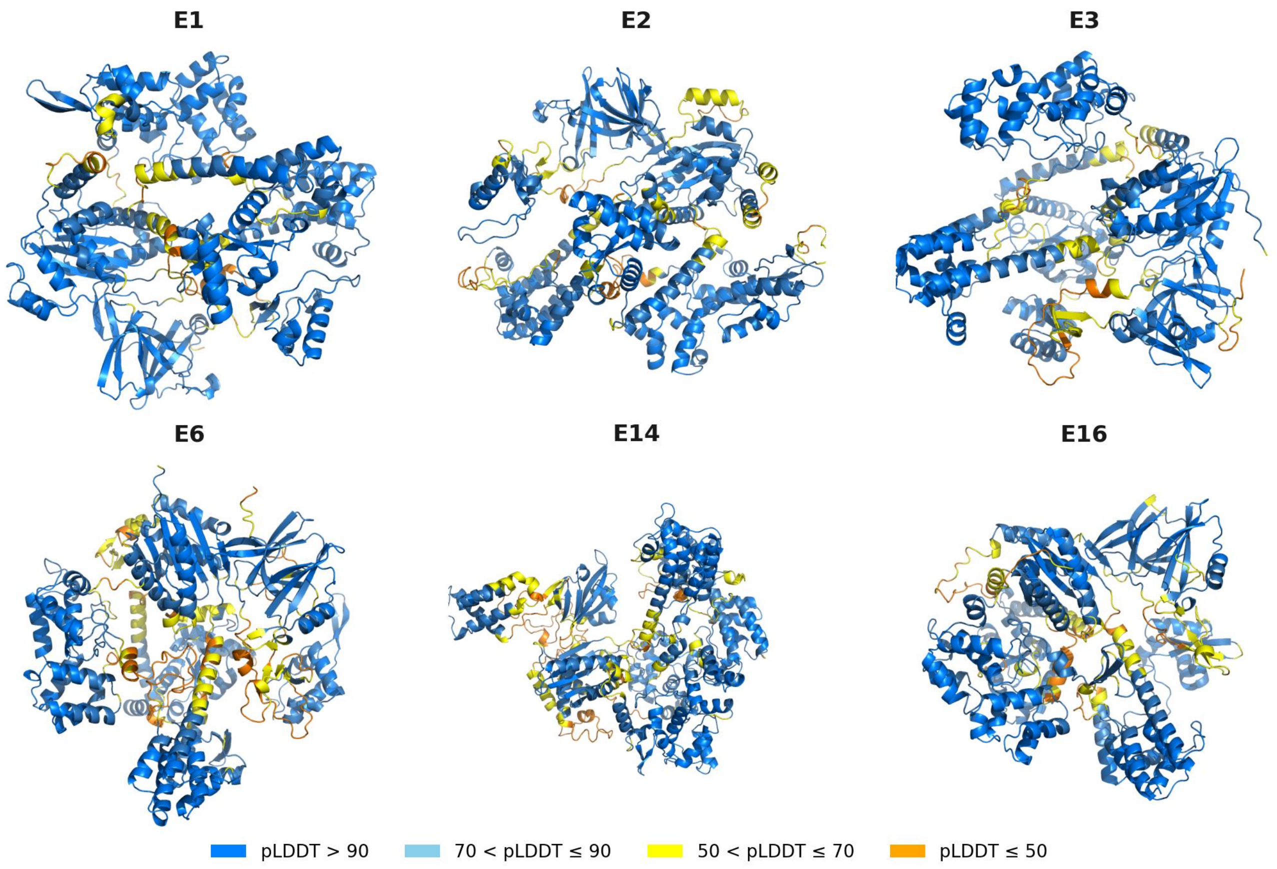
